# Transcriptional profiling reveals a previously unknown population of Cdkn2a-positive tumor-associated macrophages in aggressive brain cancer

**DOI:** 10.1101/2024.10.17.618451

**Authors:** Letícya Simone Melo dos Santos, Lucas Kich Grun, Juliete Nathali Scholl, Florencia María Barbé-Tuana, Marco Demaria

## Abstract

Microglia, the resident immune cells of the Central Nervous System (CNS), play crucial roles in homeostasis, immune responses, and the modulation of neuronal functions. In brain tumors, microglia are difficult to distinguish from infiltrating macrophages and are collectively called Tumor-Associated Macrophages (TAMs). While these cells exhibit plasticity, support tumor growth, and suppress antitumor immune responses, their identity and function remain largely unknown. Cellular senescence, a state of growth arrest associated with the upregulation of the cell cycle inhibitor p16^INK4A^ (Cdkn2a), can both suppress and promote tumor progression via the secretion of inflammatory and tissue remodeling factors, collectively known as Senescence-Associated Secretory Phenotype (SASP). By leveraging existing single-cell RNA sequencing (scRNA-seq) data of murine brain tumors, we identified a population of Cdkn2a-positive TAMs involved in immune regulation and neurogenesis, characterized by high expression of Emb and Cd93, which we named MECC (Microglia-like, and Emb-, CD93- and Cdkn2a-positive) cells. In humans, these cells were more prevalent in recurrent glioblastoma and associated with poorer prognosis. Comparative analysis of brain samples from patients with neurodegeneration and healthy older individuals revealed the specificity of MECCs for brain tumors. These findings suggest that MECCs are a unique and distinct cell population associated with aggressive brain tumors, which could potentially serve as targets for novel treatments.

## Introduction

Microglia are the resident immune cells of the central nervous system (CNS) and play a pivotal role in maintaining homeostasis, mediating immune responses, and modulating neuronal functions. In a healthy state, microglia constantly monitor the brain environment, facilitating synaptic pruning, clearing debris, and secreting neurotrophic factors that are essential for neural health (Prinz et al., 2019). Upon activation by injury, infection, or disease, microglia undergo morphological changes and shift their gene expression profiles to adopt either neuroprotective or neurotoxic phenotypes (Thompson and Tsirka 2017). Activated microglia can release proinflammatory cytokines, chemokines, and reactive oxygen species, contributing to neuroinflammation and potentially exacerbating neurological conditions (Qin et al.2023).

In brain tumors, microglia and infiltrating macrophages are collectively referred to as Tumor-Associated Macrophages (TAMs). This is because, at the transcriptomic level, distinguishing microglia from macrophages is extremely challenging owing to their overlapping profiles (Tao et al., 2023). Despite this limitation, TAMs are recognized components of the tumor microenvironment (TME) (Wang et al., 2022).

In the TME, TAMs exhibit a high degree of plasticity and display both anti- and pro-inflammatory functions (Tan et al., 2021), with an ample spectrum of intermediate populations between these two extreme states (Murray, 2017). TAMs polarized toward an anti-inflammatory phenotype contribute to tumor progression through various mechanisms. They promote angiogenesis by secreting pro-angiogenic factors, such as VEGF and TGF-β, facilitating tissue remodeling, and suppressing immune responses by secreting IL-10 and other immunosuppressive cytokines (Mantovani et al., 2017). Additionally, they can enhance tumor cell migration and invasion, creating a favorable environment for metastasis (Noy and Pollard, 2014). In contrast, TAMs can assume a pro-inflammatory phenotype that supports anti-tumor immunity. TAMs can produce pro-inflammatory cytokines (such as TNF-α and IL-12) and reactive oxygen species (ROS) and kill tumor cells either directly or via the stimulation of cytotoxic T cells (Mosser and Edwards, 2008). Furthermore, pro-inflammatory-polarized TAMs can enhance antigen presentation, helping activate adaptive immune responses against tumors (Augustine et al., 2022). Therefore, broad inhibition of TAMs could inadvertently suppress their beneficial anti-tumorigenic functions, but a strategy to specifically target pro-tumorigenic TAMS is lacking.

Cellular senescence is a state of stable and generally irreversible growth arrest that acts as a tumor suppressor by preventing the proliferation of damaged or stressed cells (Campisi, 2001). Senescent cells are characterized by the expression of cell cycle inhibitors, such as p16^Ink4a^ and p21^Cip1^, morphological changes, and secretion of a complex mix of pro-inflammatory cytokines, growth factors, and proteases, collectively known as the senescence-associated secretory phenotype (SASP) (Hernandez-Segura et al., 2018). While cellular senescence can suppress tumor initiation, SASP paradoxically promotes tumor progression by creating a pro-inflammatory environment that supports tumor growth, invasion, and metastasis (Schmitt et al., 2022).

Senescence affects both the microglial and macrophagic compartments, with senescent microglia and peripheral macrophages showing reduced phagocytic activity and altered expression of chemokines and cytokines. Senescent-like microglia accumulate in the aging brain and potentially contributes to neuroinflammation and neuronal damage (Talma et al., 2021). Similarly, senescent-like macrophages promote chronic inflammation and impair the resolution of inflammatory reactions in older organisms (Franceschi et al., 2018).

Despite the established role of cellular senescence in both microglia and macrophages, the specific contributions of senescence-associated TAMs within the brain TME, their interactions with other components of the TME, and their potential as therapeutic targets remain poorly understood.

Recent advancements in single-cell RNA sequencing (scRNA-seq) have revolutionized the ability to dissect cell heterogeneity and functional states within the TME (Zhao et al., 2023). This technology enables high-resolution characterization of cellular subpopulations, revealing distinct transcriptomic profiles, and uncovering novel cell types and states. A recent study demonstrated that removing p16-positive cells reduced brain tumor growth and improved the survival of transgenic mice (Salam et al., 2023).

Here, we employed a combination of single-cell RNA sequencing (scRNA-seq) data from mouse and human samples to identify Cdkn2a-positive populations, the transcript encoding for p16, and senescent-like cells exclusively associated with aggressive brain tumors.

## Results

### p16 elimination reduces tumor-associated microglia-like and macrophage populations

To evaluate whether p16-positive and senescent-like cells were associated with brain cancer, we used a publicly available scRNA-seq dataset of brain tumors collected from wild-type or p16-depleted mice (Salam et al., 2023). Initially, we visualized the 3MR and WT single-cell dataset distributions over diverse cell populations present in the brain tumor microenvironment (Fig. 1a). Next, we screened for Cdkn2a-positive cells expressing a well-established senescence gene list, called SENMAYO, across the tumor microenvironment (Fig. 1b and 1c). Interestingly, the Cdkn2a-positive cells with the highest SENMAYO score also expressed various TAMs markers, such as *Aif1, Tmem119, Cx3cr1, P2ry12, Aif1*, and *Csf1r* (Fig. 1d and S1a). Further analysis of TAMs cells identified two distinct clusters, 0 and 1 (Fig. 1e), which exhibited reductions of 60% and 80%, respectively, in p16-depleted mice (Fig. 1f). Because cluster 1 showed a pathological profile exemplified by the expression of disease-associated markers such as S100a10 (Afanador et al., 2014) and Mt3 (Juárez-Rebollar et al., 2017) (Fig. 1g), we focused on this subset for further analyses. In line with the sharp decline in Cdkn2a-positive cells, we observed a significant decrease in the SENMAYO score in TAM cluster 1 of p16-depleted mice (Fig. 1h), further suggesting a link between TAM cluster 1 and senescence-like state. Next, we aimed to identify surface markers that could serve as easily accessible targets on the cell membrane and further characterize the properties of TAM cluster 1. To this end, we used the EXCESP gene list, a curated set of genes encoding surface proteins (Dhusia et al., 2022). From this list, we identified Emb (Embigin) and Cd93, both of which are involved in cell adhesion, as being highly expressed in TAM cluster 1 (Fig. 1i). Cells positive for *Cdkn2a, Emb*, and *Cd93* were classified as MECCs (Microglia-like Emb-, Cdkn2a-, and Cd93-positive Cells). Notably, when considering the TAM compartment across all cell types (Fig. 1j), the number of MECCs dropped significantly from 76 cells (2.5% of total TAMs) in control mice to just 20 cells (0.63%) in p16-depleted mice (Fig. 1k). Finally, Gene Ontology (GO) analysis of MECCs revealed an overrepresentation of genes involved in cell adhesion, immune regulation, and neurogenesis, further highlighting the unique functional role of this TAM subset (Fig. 1l).

**Fig 1.**
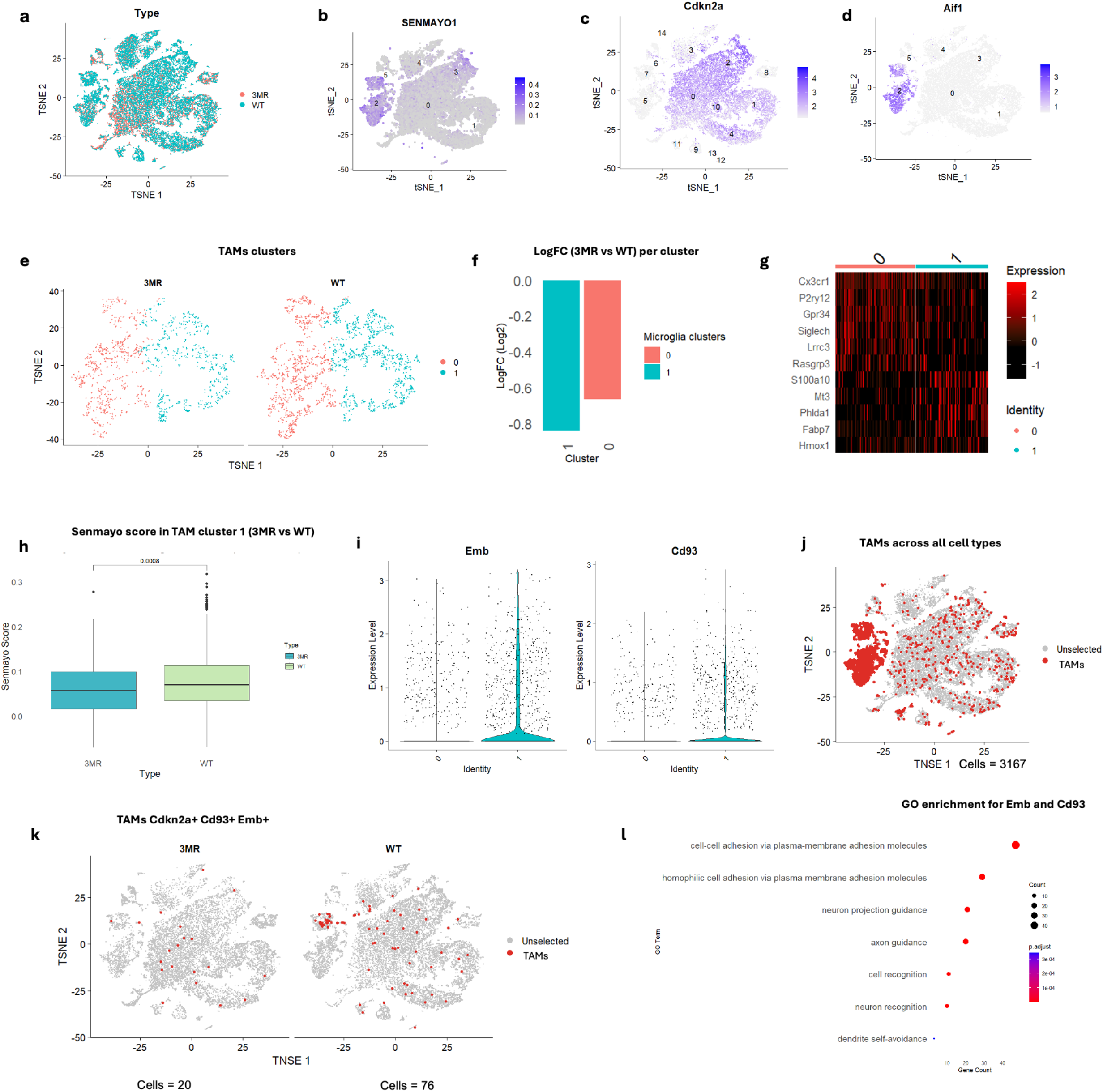
p16 Elimination reduces tumor-associated microglia-like and macrophage populations. **(a**) tSNE plot showing the distribution of cells from a scRNA-seq dataset of a glioblastoma p16-3MR mouse model from wild-type (WT) mice and mice that underwent depletion of p16-positive cells (3MR); **(b)** Representation of the SENMAYO, a list of senescence-associated transcripts, and signature expression in Cdkn2a-positive cells in the TME. **(c)** Cluster of Cdkn2a-positive cells in TME. **(d)** Evaluation of expression of the microglia-like marker Aif1 within Cdkn2a-positive cells. **(e)** Distribution of TAMs within Cdkn2a-positive clusters. **(f)** Comparison (LogFC) of Cdkn2a-positive TAM clusters between the 3MR and WT conditions. **(g)** Heat map of the expression profiles of classical and pathological TAM markers across two TAM clusters. **(h)** Analysis of SENMAYO signature expression in TAM cluster 1. **(i)** Violin plots showing Emb and Cd93 expression levels in TAM Cluster 1. **(j)** tSNE plot showing the spatial distribution of MECCs across 3MR and WT conditions. **(k)** tSNE plots illustrate MECCs (CDKN2A, EMB, and CD93) expression in 3MR mice compared to WT mice. **(l)** Gene Ontology (GO) analysis of MECCs.

### MECC’s identification and validation in human Glioma samples

We then validated the presence of MECCs in human brain tumors using an scRNA-seq dataset from glioma samples, including low-grade glioma (LGG), newly diagnosed glioblastoma (ndGBM), and recurrent glioblastoma (rGBM) (Fig. 2a). TAM cells were identified based on the expression of known markers including *AIF1, CSF1R*, and *TMEM119* (Fig. 2b). Interestingly, we observed that an increasing number of MECCs correlated with tumor aggressiveness. In LGG, we detected six MECCs (0.05%), while the number increased to 229 (2.05%) in ndGBM and 774 (6.93%) in rGBM (Fig. 2c and 2d), indicating the potential role of MECCs in tumor progression. Spatial transcriptomics further confirmed the presence of the TAM population (Fig. 2e) and their overlapping MECC signatures (Fig. 2f) in the rGBM tumor microenvironment. MECCs were identified by the overlapping expression of *CSF1R, CDKN2A, EMB*, and *CD93*. Furthermore, survival analysis of TCGA bulk RNA sequencing data revealed that high expression of Cd93 and Emb was significantly associated with reduced survival in patients with LGG (P = 0.00016) (Fig. 2g). A similar trend was observed in glioblastoma patients. However, this difference was not statistically significant (p = 0.2) (Fig. 2h). These results highlight the potential clinical relevance of MECCs in glioma progression and their association with the more aggressive forms of glioma.

**Fig 2.**
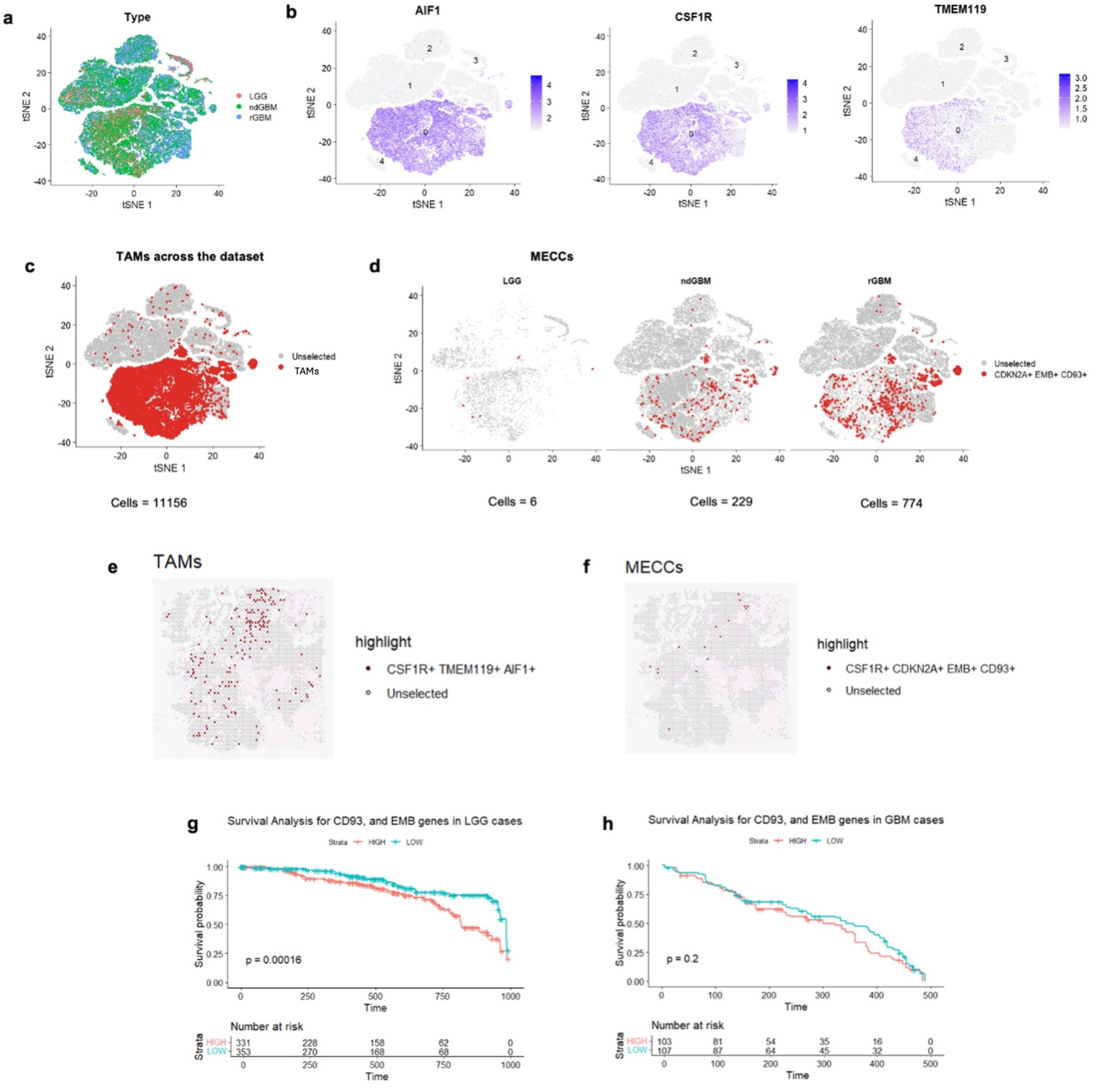
MECC’s identification and validation in human Glioma samples. **(a)** tSNE showing cell distribution in a single-cell RNA-seq dataset of human glioblastoma, including low-grade glioma (LGG), newly diagnosed glioblastoma (ndGBM), and recurrent glioblastoma (rGBM).. **(b)** tSNE plots showing TAMs AIF1, CSF1R, and TMEM119 expression across the dataset; **(c)** tSNE plot showing the selection of TAMs, based on the expression of AIF1, CSF1R, and TMEM119, established TAM markers within the dataset; **(d)** tSNE plot showing the co-expression of CDKN2A, EMB, and CD93 transcripts across the conditions (LGG – low-grade, ndGBM – newly diagnosed, and rGBM – recurrent) and its subsequent quantification; **(e)** spatial transcriptomics showing TAMs distribution, based on the CSF1R, TMEM119, and AIF1 co-expression across the tissue; **(f)** spatial transcriptomics showing CDKN2A, EMB, and CD93 co-expression location in the tissue; **(g)** and **(h)** survival probability plots for human samples with low and high expression of CD93 and EMB in LGG and glioblastoma.

### MECCs are associated with both primary and metastatic brain tumors

To explore the specificity of MECCs to the TAM cell compartment in the brain, we analyzed scRNA-seq datasets from brain metastatic melanoma (BM) and primary skin melanoma (acral-AM and cutaneous CM subtypes) (Fig. 3a and 3b). This comparative approach sought to determine whether the Cdkn2a+, Emb+, and Cd93+ signatures were associated with the brain microenvironment, but independent of tumor origin. Our analysis revealed that MECCs were almost exclusively associated with metastases, as they represented 3.86% of brain metastatic melanoma cells compared to only two 0.38% of acral melanoma cells and one 0.19% of cutaneous melanoma cells (Fig. 3c and 3d). These findings also suggest that cells expressing Cdkn2a, Emb, and Cd93 are a consequence of the brain TME, independent of whether the tumor is primary or metastatic.

**Fig 3.**
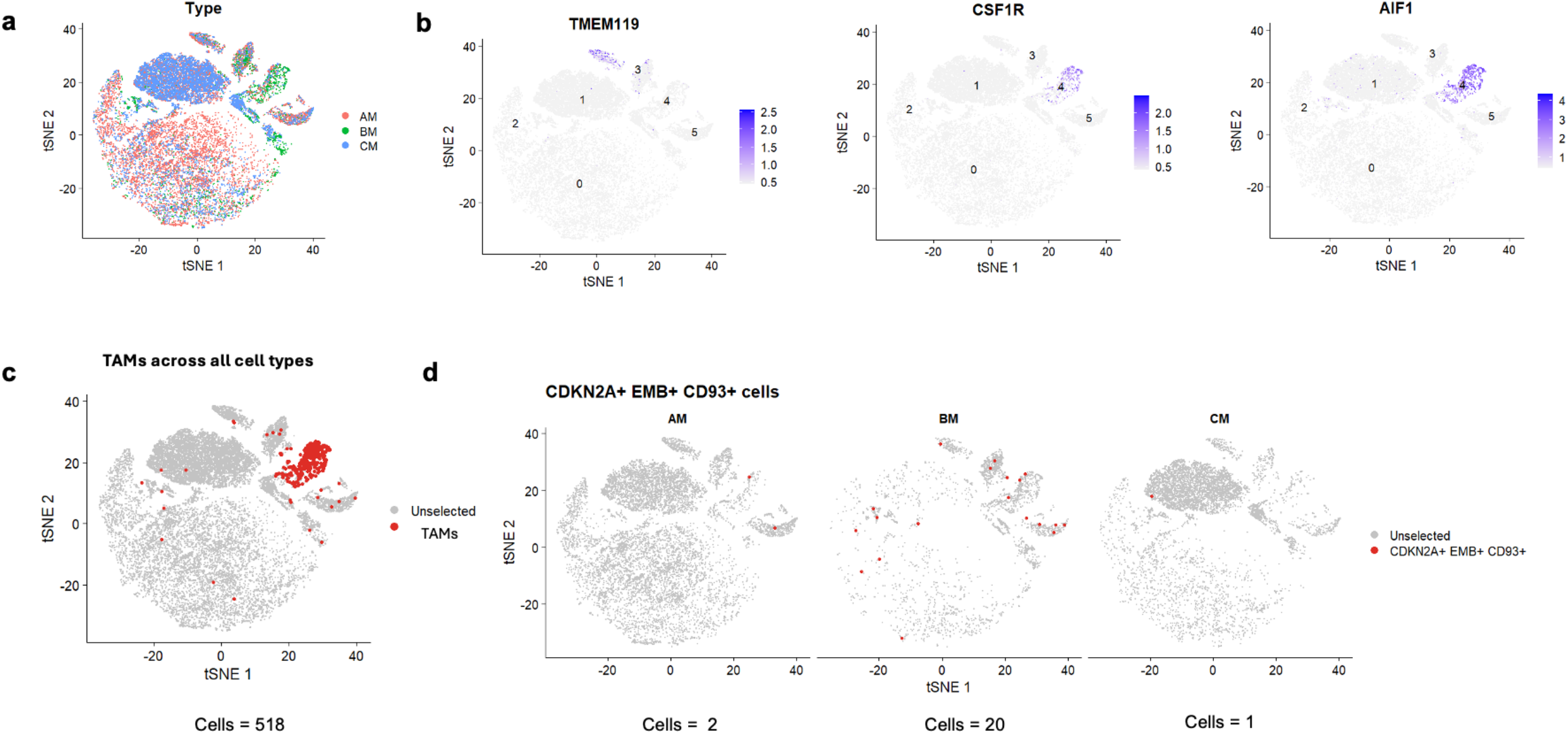
MECCs are associated with both primary and metastatic brain tumors. **(a)** Distribution of cells based on gene expression profiling of a scRNA-seq dataset of melanoma and brain metastasis datasets; **(b)** tSNE showing the TAMs AIF1 and CSF1R expression signatures across the dataset; **(c)** tSNE plot highlighting the AIF1 and CSF1R expression distribution across the dataset; **(d)** tSNE plot showing the co-expression of CDKN2A, CD93, and EMB transcripts, the MECCs population, across the conditions (AM - Acral skin Melanoma, BM, brain metastatic Melanoma, CM, cutaneous skin melanoma) and their subsequent quantification.

### MECCs are not associated with aging and neurodegeneration

Brain pathologies are significantly more prevalent in the elderly population. To explore whether the occurrence of MECCs is associated with age, we examined the TAM population in a dataset encompassing individuals aged one month to 85 years (Fig 4a and 4b). However, we did not observe a significant presence of MECCs across any age group (Fig. 4c and 4d), suggesting that aging alone does not influence the presence of these cells. The absence of MECCs in a normally aging brain indicates that their emergence is influenced by the TME rather than by general age-related changes in the brain microenvironment.

**Fig 4.**
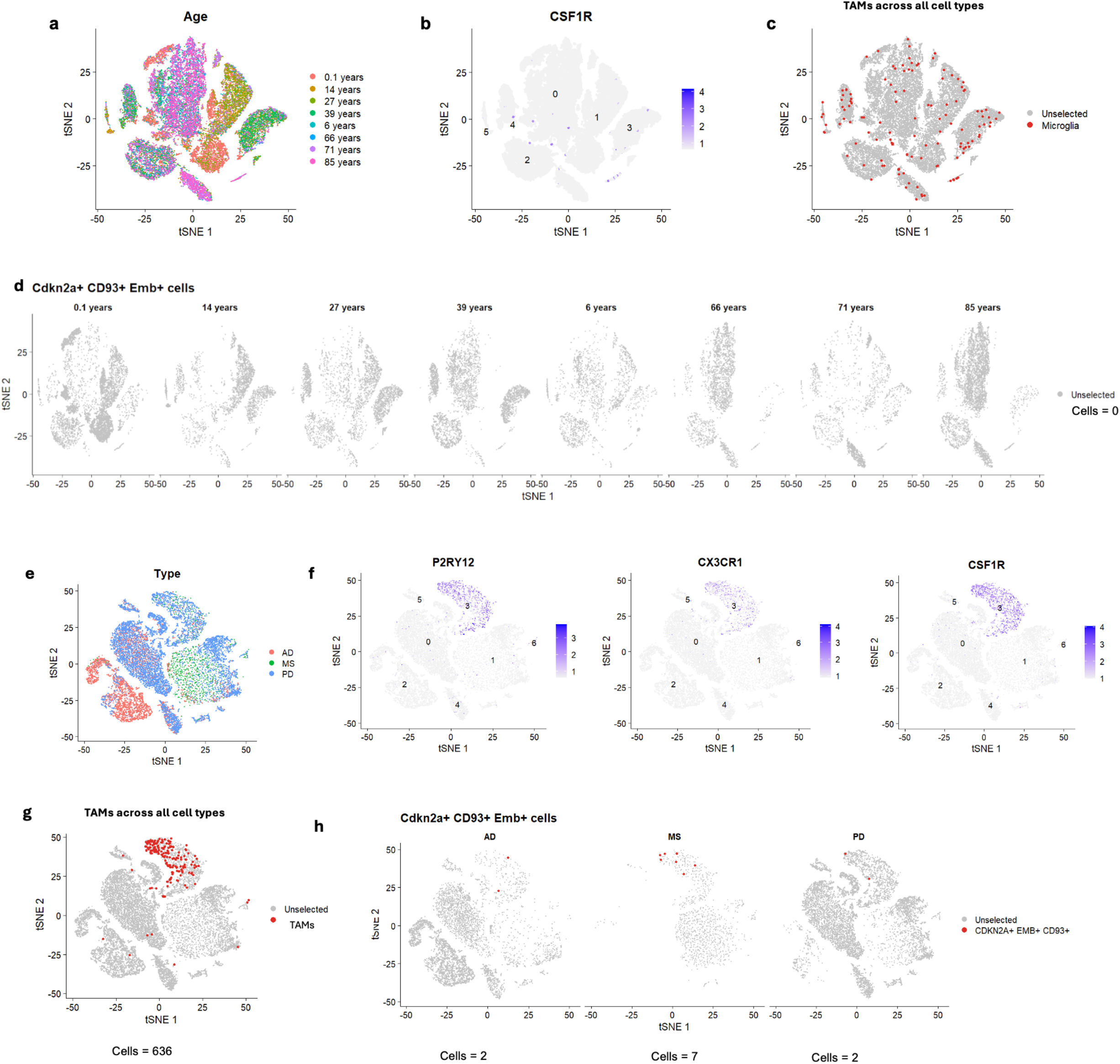
MECCs are not associated with aging and neurodegeneration. **(a)** tSNE plot illustrating the age distribution of the individuals in the scRNA-seq dataset. **(b**) Plot representing the number of CSF1R-positive cells in the TAMs across all age groups. **(c)** Spatial distribution of TAMs based on CSF1R expression across all cell types in the dataset. **(d**) tSNE plots showing the absence of MECCs signature (CDKN2A, EMB, CD93) across different age groups (0.1 years to 85 years). **(e)** tSNE plot representing cell distributions in different neurodegenerative diseases: Alzheimer’s disease (AD), Multiple Sclerosis (MS), and Parkinson’s disease (PD). **(f) The** tSNE plot showing the distribution of P2RY12, CX3CR1, and CSF1R across neurodegenerative disease conditions. **(g)** tSNE highlighting the expression of TAM markers P2RY12, CX3CR1, and CSF1R across all cell types under neurodegenerative disease conditions. **(h**) tSNE plot displaying CDKN2A, EMB, and CD93 co-expression in the AD, MS, and PD samples.

To investigate the potential general pathogenic role and pathological induction of MECCs in the brain, we assessed their presence in neurodegenerative diseases including Alzheimer’s disease, Parkinson’s disease, and Multiple Sclerosis (Fig. 4e and 4f). MECCs were identified in only two (0.31%) Alzheimer’s samples, seven (1.10%) Multiple Sclerosis samples, and two (0.31%) Parkinson’s samples (Fig. 4g and 4h). The low expression of MECCs in neurodegenerative conditions suggests that these cells are not broadly associated with all forms of brain pathologies. Instead, their presence appears to be tightly linked to cancerous conditions, particularly brain tumors and local metastases.

## Discussion

Previous research has demonstrated a strong association between senescent TAMs and poor prognosis in patients with glioma (Andersen et al., 2021). Targeting senescent cells can mitigate tumor progression and enhance therapeutic efficacy in cancer (Wang and Demaria, 2021). However, because the characterization of individual populations of senescent-like TAMs remains limited, this approach has not yet been clearly explored in brain tumors.

In this study, we identified distinct Cdkn2a-positive TAM populations in brain tumors. One of these, which we named MECC, showed the highest reduction in transgenic mice depleted of p16-positive cells and substantial senescence-like and pathological features, exemplified by the upregulation of the senescence signature SenMayo and genes involved in immune regulation and neurogenesis.

MECCs express two unique surface proteins: Cd93 and Emb. Cd93 has been identified as a receptor for C1qr, a complement protein. However, further analysis showed that Cd93 does not function in the complement system but is involved in cell-cell and cell-molecule adhesion (Galvagni et al., 2016; McGreal et al., 2002). Notably, Cd93 has been previously identified as a marker of glioma progression, highlighting the complexity of its role in the TME (Ma et al., 2022). EMB, a protein belonging to the immunoglobulin superfamily, has been poorly characterized in cancer, making it a potential new target for cancer-related conditions (Lain et al., 2009). It is essential to acknowledge that Cd93 and Emb markers individually may not be unique to TAM cells, and that their expression may be observed in other cell types, emphasizing the need for further research to elucidate their specific roles in the MECC phenotype. The increasing presence of MECCs in recurrent glioblastomas indicates their association with pathological conditions and potential as biomarkers for disease progression and therapeutic responses.

Importantly, we observed a significant correlation between Emb and Cd93 and reduced survival in patients with glioma, underscoring the clinical relevance of MECCs and their potential as novel therapeutic targets in aggressive brain tumors. It is important to emphasize the specificity of MECCs in the context of brain tumors. The higher prevalence of MECCs in aggressive primary brain tumors and metastases emphasizes the unique microenvironment of the brain, which may facilitate the development and maintenance of these cells. The extremely low representation of MECCs in other brain pathologies indicates a context-specific role for these cells in cancer and neurodegeneration. It is well known that microglia and senescent-like microglia expressing p16 and p21 markers and increased senescence-associated beta-galactosidase activity (SA-β-gal) are present in neurodegenerative conditions such as Alzheimer’s disease and Parkinson’s disease (Melo dos Santos et al., 2024). The negligible percentage of MECC cells identified in neurodegenerative conditions could indicate the heterogeneity of senescent microglia- and macrophage-like cells in the brain, suggesting the need to carefully evaluate various anti-senescence approaches tailored to specific pathologies.

Although our study provides crucial insights into the role of p16-positive TAM cells in brain tumors, several limitations warrant consideration. The reliance on animal models and retrospective analysis of human datasets may only partially capture the dynamic interactions within the tumor microenvironment. As mentioned above, the direct or indirect role of p16 elimination in these cells must be clarified, with a better understanding of whether these cells are suppressive or progressive in the tumor microenvironment. Our data suggests correlations between aggressive brain tumors and lower survival probabilities, but additional tests are needed to determine the real impact of MECCs on the TME in GBM. Future research should address these limitations by incorporating advanced imaging techniques, in vitro co-culture models, induction of a senescent-like state, and longitudinal patient studies to better understand the functional impact of MECCs in aggressive brain tumors. Another limitation is that although Emb and Cd93 transcripts are membrane proteins, we did not check their proteomic expression, so further analysis should validate their presence through techniques such as flow cytometry and immunofluorescence. At the protein level, while Cd93 can be identified on the cell surface, whether it can be co-expressed with Emb remains an open question that needs to be further investigated. Additionally, our study focused on microglial cells, but it is well-established that microglia and monocyte-derived macrophages (MDMs) are difficult to distinguish in the TME due to their transcriptomic similarities. We used a nomenclature that comprehensively embraces both populations, but future studies should address whether we should continue treating them as one population or different populations, what are their functional and phenotypic differences, and their contribution to the MECC population. Elucidating the specific roles of microglia and MDMs in the MECC phenotype is essential for developing targeted therapies that modulate senescent-like TAMs in brain tumors.

In conclusion, our study revealed the significant contribution of p16-positive TAM cells, specifically MECCs, to the brain TME. We highlighted their potential as therapeutic targets by identifying the transcriptomic profiles of these cells and their correlation with aggressive brain tumors. Further research on the mechanistic underpinnings and clinical validation of MECC-targeted therapies is warranted to advance treatment strategies for patients with brain tumors. These findings underscore the urgent need for targeted therapies to modulate senescent-like TAMs in brain tumors, potentially transforming treatment paradigms and improving patient outcomes.

## Methods

### scRNAseq analysis

We analyzed single-cell RNA sequencing (scRNA-seq) data using Seurat and other relevant packages in R. The analysis involved several key steps including data preprocessing, normalization, clustering, and differential expression analysis. Each dataset was imported using either the ReadMtx function, generating the expression matrix from the raw data, using Read10X_h5 or read_rds, and converted into a Seurat object using CreateSeuratObject. The metadata for each sample were annotated, including sample information and sample type.

### Quality Control and Filtering

The mitochondrial gene percentage was calculated using PercentageFeatureSet. Cells with more than 5% mitochondrial content or less than 800 Unique Molecular Identifiers (UMIs) and 500 detected genes were filtered out.

### Data Integration

Individual Seurat objects are merged into a single Seurat object. The merged object underwent normalization using the NormalizeData function, and the variable features were identified using FindVariableFeatures. The data were scaled using ScaleData and principal component analysis (PCA) was performed using RunPCA. To address the batch effects, we integrated the data using the harmony integration pipeline. After the integration, the data were scaled again. We performed a nearest neighbor analysis using FindNeighbors and clustering using FindClusters. t-distributed Stochastic Neighbor Embedding (t-SNE), a popular technique for dimensionality reduction in large datasets, was used for dimensionality reduction and visualization (RunTSNE).

### Differential Expression Analysis

We identified differentially expressed genes using FindAllMarkers, focusing on genes with a minimum percentage threshold of 0.25 and a log fold change threshold of 0.25. Specific genes and clusters of interest were further analyzed, and the top marker genes for each cluster were identified.

### Gene Set Enrichment Analysis

Gene ontology (GO) enrichment analysis was performed using the clusterProfiler package.

The enriched biological processes were visualized using dot plots.

### Visualization

Various plots were generated to visualize the data and results, including violin plots (VlnPlot), feature plots (FeaturePlot), and heatmaps (DoHeatmap). Differential expression results were visualized using Enhanced Volcano for specific clusters and conditions. Specific cells expressing genes such as CDKN2A, EMB, and CD93 were highlighted using WhichCells and DimPlots.

### Spatial Transcriptomics Analysis

The spatial transcriptomic dataset was loaded using the read_rds function. Data normalization was performed using the NormalizeData function in a spatial assay. The percentage of mitochondrial genes is calculated with PercentageFeatureSet using the pattern ^MT-. Cells were filtered to retain those with more than 800 UMIs, at least 200 detected features, and less than 5% mitochondrial content. Variable features were identified using FindVariableFeatures. Data was scaled using ScaleData. PCA was performed with ‘RunPCA.’ Like scRNA-seq, nearest neighbors were found using FindNeighbors, and clustering was performed with FindClusters. Dimensionality reduction for visualization was achieved using RunTSNE. Markers were identified using FindAllMarkers. Spatial plots were created using SpatialFeaturePlot to visualize the spatial distribution of genes, such as TMEM119, CX3CR1, P2RY12, AIF1, CSF1R, ITGAM, CD93, EMB, and CDKN2A. Cells expressing specific gene combinations (e.g., CSF1R, CDKN2A, EMB, CD93) were identified using ‘WhichCells‘ and highlighted with ‘SpatialDimPlot.’

### The Cancer Genome Atlas Program (TCGA) Analysis

The clinical data for the TCGA-LGG cohort was retrieved using the GDCquery_clinic function. Relevant columns for survival analysis were extracted, and vitalstatus was encoded as deceased status. Overall_survival was calculated based on days_to_death for deceased patients and days_to_last_follow_up for living patients. A project summary for TCGA-LGG and TCGA-GBM cohorts was retrieved, and gene expression data were queried and downloaded using the GDCquery and GDCdownload functions. The data were prepared using the GDCprepare function and transformed into a matrix. The gene and sample metadata were extracted from the prepared data. The data were transformed using the vst function in DESeq2, with genes with fewer than 10 reads across all samples removed. Data for the EMB and CD93 genes were extracted, and median values were calculated. The data were stratified based on expression levels, with samples having counts greater than or equal to the mean value classified as “high” and those with counts less than the mean value classified as “low.”

### Survival Analysis

The data for EMB and CD93 were combined. Survival curves were fitted using the survfit function, the Kaplan-Meier method, and visualized with a ggsurvplot.

## Data Availability

The datasets used in this study are publicly available and can be accessed through the following repositories: NCBI GEO: GSE168038 (p16-3MR mouse model), GSE182109 (human Glioma samples), GSE186344 (human Melanoma brain metastasis), GSE215121 (human Melanoma skin subtypes), GSE146639 (human Alzheimer’s Disease), GSE140231 (human Parkinson’s Disease), GSE179427 (human Multiple Sclerosis), GSE199243 (human lifespan).

Spatial Transcriptomics: Human Glioblastoma Whole Transcriptome Analysis (https://www.10xgenomics.com/datasets/human-glioblastoma-whole-transcriptome-analysis-1-standard-1-2-0)

The Cancer Genome Atlas (TCGA): https://portal.gdc.cancer.gov

