## Supplementary Figure 1 for "Transcriptional profiling reveals a previously unknown population of Cdkn2a-positive tumor-associated macrophages in aggressive brain cancer"

**Extended Data**

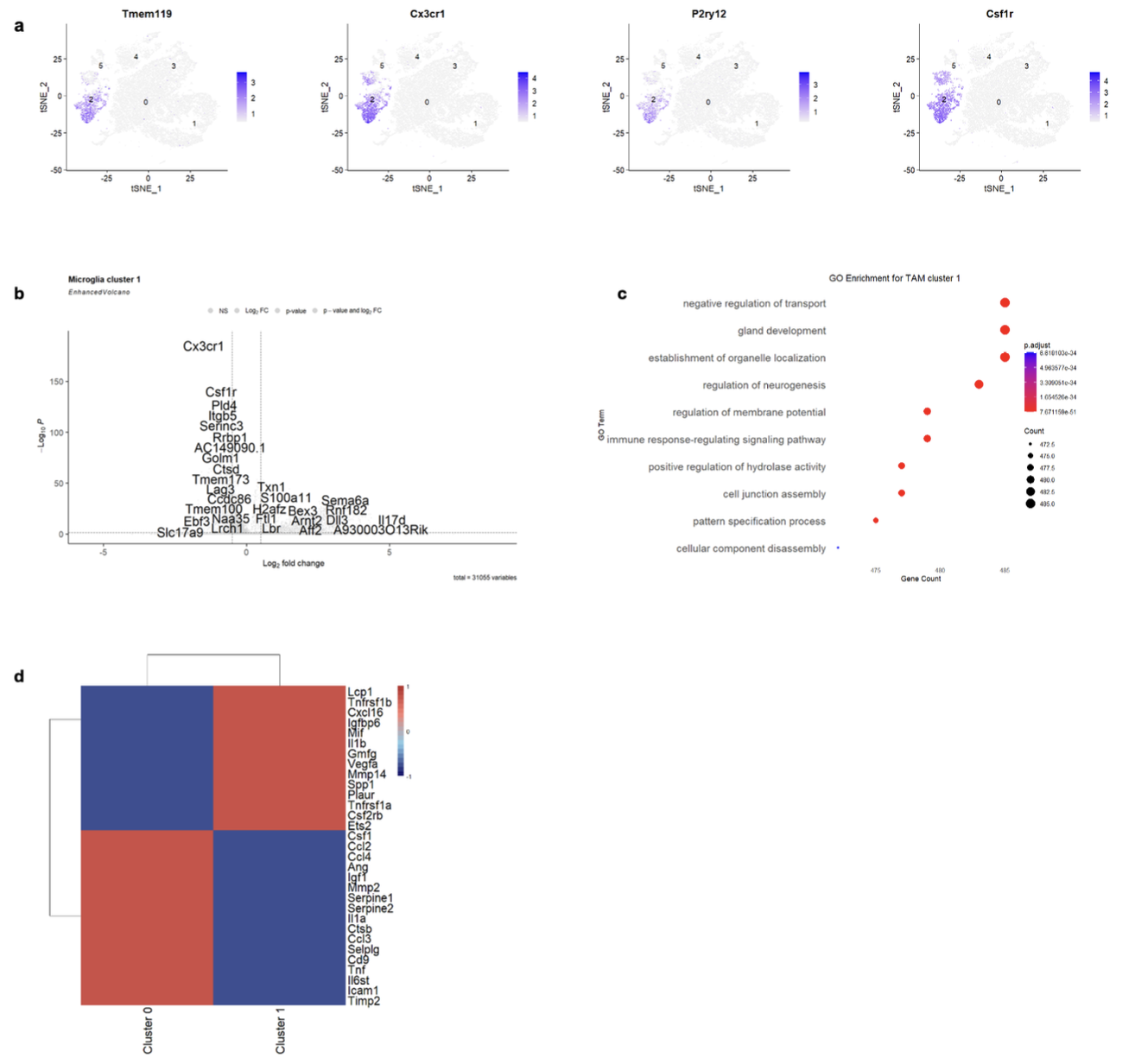

***Fig. S1. Characterization of TAM cluster 1 and gene expression profiles in TAMs***
***(a)*** *Expression patterns of canonical microglia markers, including Tmem119, Cx3cr1, P2ry12, and Csf1r, across TAM populations in the tumor microenvironment. These markers confirm the identification of TAMs as microglia-like cells within the tumor.* ***(b)*** *Volcano plot highlighting differentially expressed genes in TAM cluster 1.* ***(c)*** *Gene Ontology (GO) enrichment analysis for TAM cluster 1, reveals enrichment for biological processes such as the regulation of neurogenesis, immune signaling, cell junction assembly, and cellular disassembly.* ***(d)*** *Heatmap comparing the SENMAYO gene expression profiles of two TAM clusters, cluster 0 and cluster 1.*
